# Maintaining tiger connectivity and minimizing extinction into the next century: Insights from landscape genetics and spatially-explicit simulations

**DOI:** 10.1101/081117

**Authors:** Prachi Thatte, Aditya Joshi, Srinivas Vaidyanathan, Erin Landguth, Uma Ramakrishnan

**Author notes:** Corresponding authors; National Centre for Biological Sciences, Tata Institute of Fundamental Research, Bellary Road, Bangalore 560065. /31.

## Abstract

Habitat loss is the greatest threat to large carnivores around the world. Maintenance of functional connectivity in fragmented landscapes will be important for long-term species persistence. Here, we merge landscape genetics analyses and spatially-explicit simulations to understand future persistence and extinction of tigers (*Panthera tigris*) in Central India. Tigers in this landscape are restricted to Protected Areas (PAs) and forest fragments embedded within a mosaic of agricultural fields and human settlements. We examined current population connectivity of tigers across nine reserves (using 116 non-invasively sampled individuals and 12 microsatellites). Genetic data was used to infer resistance-to-movement. Our results suggest that dense human settlements and roads with high traffic are detrimental to tiger movement. We used landscape genetic simulations to model 86 different scenarios that incorporated impacts of future land-use change on inferred population connectivity and extinction. Our results confirm that genetic variability (heterozygosity) will decrease in the future and small and/or isolated PAs will have a high risk of local extinction. The average extinction risk of small PAs reduced by 23-70% on adding a 5 km buffer around exiting boundaries. Unplanned development results in 35% lower heterozygosity and 56% higher average extinction probability for tigers even within protected areas. Increasing tiger numbers in such a scenario decreases extinction probability just by 12 % (from 56% to 44%). Scenarios where habitat connectivity was enhanced and maintained, stepping-stone populations were introduced/maintained, and tiger numbers were increased, led to low overall extinction probability (between 3-21%). Our simulations provide a means to quantitatively evaluate the effects of different land-use change scenarios on connectivity and extinction, linking basic science to land-use change policy and planned infrastructure development.

## Introduction

The current rates and magnitude of species decline and extinction are higher than ever before (Barnosky *et al*. 2011; Dirzo *et al*. 2014). Most mammals retain less than half of their historical range, resulting in substantial population decline and habitat fragmentation (Morrison *et al*. 2007; Dirzo *et al*. 2014). Due to their large area requirements, slower life histories and low densities, large carnivores are especially vulnerable to habitat fragmentation and isolation. A majority (77%) of large carnivores continue to undergo worldwide decline, with populations at risk of local extirpation due to habitat loss (Ripple *et al*. 2014).

Conservation efforts, including population monitoring, legal protection, creation of protected areas, reintroductions and translocations, have ensured recovery of species, such as the grey wolf (*Canis lupus*) in North America and brown bears (*Ursus arctus*) in northern Europe, among others (Ripple & Beschta 2012; Hagen *et al*. 2015). Long-term persistence of such threatened populations requires identifying and maintaining connectivity among habitat patches (Jackson *et al*. 2016). Among large carnivores, significant attention and resources are invested in recovery and conservation of the tiger, an iconic species with less than 4000 individuals left in the wild.

Tigers have lost four subspecies and 93% of their historical range, and what remains of their existing range is highly fragmented. With nearly 65% of the world's wild tigers (Jhala, Y. V.;Qureshi, Q.;Gopal 2015) and substantial genetic variation (Mondol *et al*. 2009a), India is a stronghold for tiger survival. Recent reports suggest that conservation and management efforts in India over the last three decades have led to a 30% increase in tiger numbers (Jhala, Y. V.;Qureshi, Q.;Gopal 2015). Despite what appears to be a demographic recovery, the median number of tigers within individual protected areas (PA) in India is low (median: 19, range: 2-215; (Wikramanayake *et al*. 2010; Jhala, Y. V.;Qureshi, Q.;Gopal 2015)). Most populations by themselves may not be viable, and the continued survival of tigers could be contingent on maintaining connectivity between PAs.

Several independent genetic studies in the high priority tiger conservation landscape of Central India confirm that PAs exchange dispersing individuals and are fairly well connected (Joshi *et al*. 2013; Sharma *et al*. 2013a; Yumnam *et al*. 2014; Reddy *et al*. 2017). About 35% of India's tigers are estimated to live outside PAs (Jhala, Y. V.;Qureshi, Q.;Gopal 2015) and may play a critical role in maintaining connectivity. However, India is a country with over a billion people, an economy growing at 7% annually, poised for rapid urbanization and the ensuing increase in associated infrastructure. Among the planned infrastructure, existing highways in the landscape are being widened to meet the demands of increasing traffic (e.g., National Highway 7 which bisects a critical corridor is being widened after a prolonged legal battle owing to the conflict between tiger conservation and development activities (Srivastava & Tyagi 2016)), further fracturing an already highly fragmented landscape. Landscapes outside PAs (~95% of India’s area) are about to change dramatically, and tiger management and conservation is currently only focused on protected areas.

Earlier studies correlating genetic connectivity with landscape elements have revealed that tiger movement is negatively impacted by human settlements within the Central Indian landscape (CIL) (Joshi *et al*. 2013). Urban populations in India are projected to double from 410 million in 2014 to 814 million in 2050 (United Nations *et al*. 2014). Additionally, built-up areas have been increasing almost 3 times faster than population in nearly all large Indian cities (Sudhira 2011). Along with the urbanization, demands for better road and railway connectivity between cities is also projected to increase (National Transport Policy Development Committee 2013). Such landscape transitions will negatively impact tiger connectivity. Small, isolated populations have known genetic consequences, including low variation (Frankham 1996; Boersen *et al*. 2003), increased inbreeding and increased disease susceptibility (Spielman *et al*. 2004; Trinkel *et al*. 2011), and heightened extinction risk (Saccheri *et al*. 1998). However, these are general predictions, and we do not know how future land-use change will specifically impact connectivity and persistence of tigers in the landscape.

Few studies (Tian *et al*. 2011, 2014) have attempted to understand how future climate and landscape change might affect persistence of tigers. Tian *et al*. (2014) used population viability analysis (PVA) to simulate the effect of climate change and habitat fragmentation on future persistence of Amur tigers by incorporating factors affecting species distribution, but not dispersal. Persistence of tigers in complex and changing landscapes such as CIL requires modeling population persistence based on empirical genetic data, factors affecting dispersal, predicted landscape/climate change and interactions of these factors with demography. Globally, very few such intensive modeling exercises have been conduct to predict future persistence of endangered species (Landguth *et al*. 2014; Benson *et al*. 2016; Brown *et al*. 2016).

In this paper, we examine genetic connectivity among tiger populations in the CIL, including individuals within and outside PAs. We use this data on gene flow to infer the effect of different landscape features on dispersal and connectivity. We then carry out forward-time, spatially-explicit, individual-based simulations to understand how genetic diversity, connectivity and extinction probability will change under nine different development scenarios. We examine these scenarios, while accounting for tigers inside and outside PAs. We also test the effect of increasing tiger numbers and the effects of assumptions about dispersal, modeling a total of 86 scenarios.

## Materials and Methods

In this study, we collected genetic samples from wild populations (*Study area and sampling*), generated genetic data and conducted population genetic analyses (*Genotyping and population genetic analysis*). Landscape genetic analyses allowed us to infer landscape elements impacting connectivity (*Landscape genetic analyses*). Future scenarios were simulated assuming various criteria for landscape change, including specific management relevant scenarios, and tiger demography (*Landscape genetic simulations*). All simulations inferred genetic variability, inbreeding, connectivity and extinction probability in 2100. A flow chart of the methods is presented in Figure S1 and more detail is described as follows.

### Study area and sampling

The CIL is a global priority tiger conservation landscape. With ~34% forest cover and an estimated 688 (596-780) tigers (Jhala, Y. V.;Qureshi, Q.;Gopal 2015), it is a stronghold for tiger conservation. The PAs in the landscape are embedded in a heterogeneous matrix of multiple land-use types.

Non-invasive (scat) samples (n= 580) were collected between October 2012 and April 2014 from potential areas (PAs and forested areas outside PAs which are a part of territorial forest divisions and forest development corporations) in the state of Maharashtra, Madhya Pradesh and Chhattisgarh. We sampled eleven PAs: (1) Kanha Tiger Reserve (KTR), (2) Pench Tiger reserve (PTR), (3) Bandhavgarh Tiger Reserve (BTR), (4) Achanakmar Tiger Reserve (ATR), (5) Nagzira (NGZ) and (6) Nawegaon (NAW) (which together comprise a Tiger Reserve) (7) Satpura Tiger Reserve (STR), (8) Tadoba-Andhari Tiger Reserve (TATR), (9) Bor Tiger Reserve (BOR), (10) Umred-Karhandla Wildlife Sanctuary (UK) and (11) Tipeshwar Wildlife Sanctuary (TIP). We also sampled in three territorial forest divisions outside PAs: (1) Balaghat Forest Division (BAL), (2) Central-Chanda Forest Division (CHA) and (3) Bramhapuri Forest Division (BPR) (Figure 1). Sampling was also carried out in Sitanadi-Udanti Tiger Reserve (S-U) and Panna Tiger Reserve (PAN). We did not find any tiger scat samples in S-U and samples from PAN were not used to optimize the resistance layers since tigers in PAN have been reintroduced from KTR and PTR after they went extinct locally in 2006. However, we included PAN in the forward-time simulations. See supplementary material S1 for more details.

**Figure 1.**
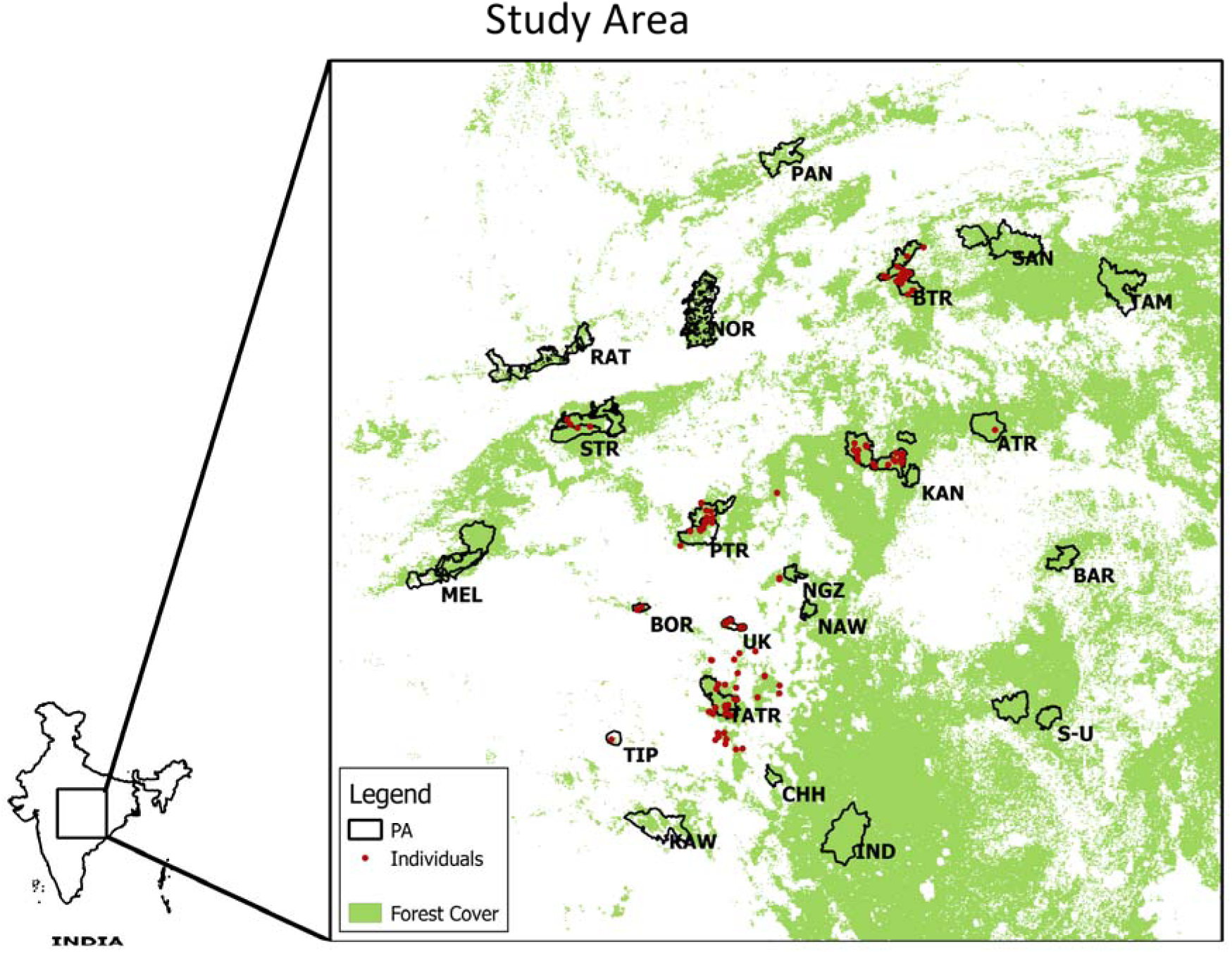
Study area: The figure contains a map of India with the study landscape highlighted and enlarged. In the enlarged study area, protected areas are marked by black outline and genetically identified individual tigers as red dots.

### Genotyping and population genetic analysis

In order to quantify genetic connectivity between different PAs, we first extracted DNA using standardized methods and identified individuals(Mukherjee *et al*. 2007; Mondol *et al*. 2009b). We then calculated heterozygosity based differentiation statistics (PopGenReport (Adamack & Gruber 2014), MMOD (Winter 2012), and HIERFSTAT (Goudet 2005) in R (Ihaka & Gentleman 2012)). For further details about genetic analysis, refer to the supplementary material S2.

### Landscape genetics analysis

We calculated inter-PA genetic distances based on proportion of shared alleles (D_PS_; (Bowcock *et al*. 1994)) using Microsatellite Analyzer (MSA, version 4.05; (Dieringer & Schlutterer 2003)). The relationship between the observed genetic structure and the landscape variables likely to affect tiger dispersal was systematically evaluated using a multi-model inference and optimization approach (Shirk *et al*. 2010) described next.

We selected landscape variables known to affect tiger dispersal based on Joshi *et al*. (2013) (i.e., land cover, human settlement layer, roads and railway lines along with density of linear features) to build resistance models. We did not use topography related variables as used in Reddy *et al*. (2017). Tigers are habitat generalists (ranging from 0-3000 meters above sea level), their occurrence is associated with forests in the CIL and they have been reported to avoid desserts and short grasslands (Kitchener and Dugmore 2000, Yumnam et al. 2014). Athreya *et al*. (2014) revealed that tigers disperse through and use less rugged areas in Central India. Additionally, over the last 300 years most forest cover loss has been in the low elevation and less rugged areas. As a result, the current forest cover is concentrated in highly rugged and relatively high-slope areas ((Sharma *et al*. 2013b), supplementary figure S3).We used MODIS land-cover data (MCD12Q1) and reclassified it into five broad land-cover types which we ranked in order of increasing resistance: (i) forest, (ii) degraded and scrub forest, (iii) agriculture (including fallow/ wasteland) and (iv) built-up (developed with buildings and non-building structures) areas. We developed a layer of human settlements by merging urban and peri-urban areas derived from nightlight data (available at National Geophysical Data Centre) and rural areas from population density data (available at http://www.worldpop.org.uk/data/get_data/). Since all villages in the study landscape are not electrified, using nightlights alone would have underrepresented human settlements within the landscape. A vector layer of national highways, state highways and major public roads was reclassified into 5 categories based on the intensity of traffic on the road (based on Passenger Car Unit (PCU) data for 2006 from the Ministry of Road Transport and Highways). PCU represents the number of vehicles in terms of passenger cars and accounts for all types of vehicles. See supplementary material S3 for more details. Railway lines and roads were used to generate a layer representing the density of linear features.

Landscape resistance values were inferred from the genetic data using multi-model optimization approach described in Shirk et al (2010). Each landscape variable was related to landscape resistance using a mathematical model (supplementary material S4). Using genetic data as the response variable and systematically varying the model parameters (maximum resistance and a shape parameter to account for the relationship of the variable with resistance), we identified the best fitting model parameters for each variable. The best fitting model was identified as the one with the most significant correlation with genetic data, after controlling for the effect of geographic distance using partial Mantel tests (Smouse *et al*. 1986). The landscape variables that explained significant variation after controlling for geographic distance in univariate models were retained for further analysis. The retained variables were combined (additively) and optimized again in a multivariate context to account for interactions between different landscape variables(Shirk *et al*. 2010). We then used the estimated landscape resistance data to understand how future landscape change may alter the resistance surface (and therefore, connectivity) as described below.

### Landscape genetic simulations

We used CDPOP (Landguth & Cushman 2010) to simulate 20 non-overlapping generations (~100 years for tigers) of mating and dispersal among individuals for each scenario (scenarios described in next section). CDPOP is a spatially-explicit, individual-based simulator of population genetic processes. It simulates mating and dispersal in a finite population assigned to fixed locations, recording alleles of all individuals every generation. We initialized individuals in the simulations as being characterized by 12 loci with at most 11 alleles per locus. For the PAs with genetic data, individuals were assigned alleles based on the allele frequency distribution for that PA. For PAs without genetic data, we assigned allele frequencies based on STRUCTURE results (Figure S5) or using other information (see S5(a) for details). Individuals were initially seeded on the landscape based on the current tiger estimates (Table S4). In each generation, females gave birth to offspring (number based on a normal distribution with mean 3±2) with an equal sex ratio at birth. Adults died, and the vacant locations were occupied by dispersing individuals. Probability governing mating and dispersal related movement was based on proximity of individual locations specified by the pairwise cost-distance matrices between individuals (calculated using ‘gdistance’ in R), which were scenario-specific (see next section). A negative exponential function (Sutherland *et al*. 2000) with median dispersal distance of 85 km (Bowman *et al*. 2002) and two different maximum dispersal distances of 300km (Patil *et al*.

2011) and 500km (Bowman *et al*. 2002; Natesh *et al*. 2017) was used to model natal dispersal. If all locations were occupied, any remaining offspring not assigned to a location were eliminated (Balloux 2001).The probabilistic functions governing birth rate, mating and dispersal introduced demographic stochasticity in the model. See Table S1 for more details on simulation parameters. At the end of each simulation, we calculated genetic diversity indices (heterozygosity, inbreeding and allelic richness) and differentiation indices (Global and pairwise Fst, G’st, Jost D) (Jombart 2008; Adamack & Gruber 2014). Probability of tiger extinction for each PA was the number of times the tiger population within the PA went extinct out of 100 replicate simulation runs. We calculated sex ratio skew and fluctuations in population size across generations.

### Simulation scenarios

In order to explore how future landscape change may affect dispersal and connectivity of tiger populations, we carried out spatially-explicit genetic simulations under different landscape change scenarios. All scenarios we present are plausible, and derived from covariates which are known to influence land-use land-cover change. The objective of these scenarios was to identify the kind of landscape-wide, land-use changes that will facilitate or impede tiger connectivity, and not to identify where land-use change will occur in the future. Scenarios of landscape change were developed using Land Change Modeler (LCM) in IDRISI (SELVA, http://www.clarklabs.org) based on the change in land-use land-cover (LULC) from a combination of land cover (MODIS) and human settlements data and road expansion from 2001 to 2012. Future LULC maps were generated for 2020, 2040, 2060, 2080. See supplementary S3a for details.

Simulations to assess change in genetic connectivity in the future were carried out for nine landscape change scenarios (labelled F1-F9, see Table 1 for the scenario description and rationale). For each of these scenarios, the landscape was parameterized using optimized resistance values based on the landscape genetic analysis described in the previous section. Cost-distance matrices were calculated for all the future maps that were generated. During simulations, the distance matrices changed after every 4 generations (~20 years) to account for the changing landscape except for scenario F1 (no landscape change) and F9 (PAs were fenced at the 1^st^ time step of 20 years and remained fenced after that). Along with scenarios F1 to F9, we carried out additional simulations for scenarios F1 and F4 where we added a 5km buffer around the smaller PAs in the landscape (< 400km^2^). We evaluated the effect of increased PA size on connectivity and extinction estimates. Simulations for the first eight scenarios were carried out under 4 sub-scenarios:(a) with tigers restricted to PAs (current numbers constant), (b) with tigers inside and outside PAs (outside individuals distributed randomly within existing forest patches), (c) with tigers inside and outside PAs (outside individuals clustered in space to form ‘stepping-stone’ populations between PAs), and (d) with tigers restricted to PAs (numbers increase). See supplementary material S5b for details.

**Table 1.** Landscape change scenarios for forward time simulations.

| Scenario | Description | Rationale |
| --- | --- | --- |
| F1- No landscape change* | Status quo | Null scenario |
| F2- Forest cover constant | Landscape change modeled while keeping the forest cover constant | The Green India mission under the National Action Plan on Climate Change (Ministry of Environment and Forests 2008) advocates achieving a forest cover of 33%. Current forest cover is 21% (Forest Survey of India 2015). In 1996, the supreme court of India redefined the scope of Forest Conservation Act 1980 and banned tree felling inside forests across India (Lele & Rosencranz 2008). |
| F3- Agriculture area constant | Landscape change modeled while keeping the area under agriculture constant | Global food demand is projected to double by 2050. Even if use of technology to intensify agriculture increases yield, area under agriculture is not expected to reduce in the future(Laurance & Balmford 2013) |
| F4- Unrestricted change* | Landscape change modeled based on change from 2000 to 2012 | India's Gross Domestic Product (GDP) growth rate was higher than ever before in the decade from 2001- 2011. Although this rate reduced after 2011, the recent government's development driven policies are likely to increase the economic growth rate (Dooley *et al.* 2014). The rate of granting forest clearances has also been highest from 2002- 2011 within the last three decades. 387952 hectare of forest land was |
|  |  | diverted during this decade for defense, mining, irrigation, power projects, industries and infrastructure projects (Centre for Science and Environment 2012) |
| F5- Effect of mines and associated landuse change | In order to evaluate the effect of mines, we let the rest of the landscape remain constant (like in F1) except for the increase in area of mines and associated built-up area over the next 100 years. The mining area (mine+ built-up) increased 3.6 times every 20 years based on a study in central India (Areendran *et al.* 2013) | The central Indian region is rich in mineral deposits. The mining sector currently contributes ~2% to India's GDP and the Ministry of Mines, Government of India has targeted to increase this share to 5% of GDP (Ministry of Mines 2011). The Government of India amended the Mines and Minerals (Development and Regulation) Act in 2015 in order to expedite environmental clearances and issuance of licenses. This amendment also provides for the creation of District Mineral Foundations (DMF) to work towards developing mining affected areas. Research in Central India has shown that mining leads to landuse change and an increase in built-up areas around mines (Areendran *et al.* 2013; Prakash & Gupta 2016) and the setting up of DMF will only increase the rate of this conversion. Data on mines were obtained from the Ministry of Mines database (http://mcas.nic.in/Mining_plan_web_Query.asp) and Greenpeace India (Fernandes 2012). The data from Ministry of Mines were converted into a spatial layer based on their GPS locations and area of the mines. Data on coal mines described in (Fernandes 2012) was obtained as shapefiles through personal communication. |
| F6- Highways as barriers | Landscape does not change except two national highways (NH6 and NH7) which cut across the landscape are converted into barriers to movement | Road traffic is estimated to grow at about 13 % per annum over the next 20 years (National Transport Policy Development Committee 2013). NH7, which runs north to south, bisects a critical corridor in the landscape and has recently been cleared to be widened from two to four lane capacity. NH6, which runs east to west and bisects another critical corridor, is also being considered for widening. Yadav *et al.* (2012) have observed agriculture and built-up area encroachment along NH6 in the forested area which connects two PAs (NGZ and NAW) (Yadav *et al.* 2012), thus potentially increasing the resistance to movement of tigers. This scenario is |
|  |  | a case study to specifically look at the effects of these highways, if they were to become barriers in the future, on the corridors they bisect. |
| F7- Highways as barriers with wildlife crossings | Landscape does not change except two national highways (NH6 and NH7) which cut across the landscape are converted into barriers with provision for wildlife crossing at points where they bisect critical corridors. There was one gap in each highway. The gaps were 1 pixel wide and were placed near the locations where the NH6 and NH7 are currently narrow (2lanes wide, without a divider in between) and there is forest cover on the either side. | Guidelines to mitigate impacts of linear infrastructure on wildlife recommend a 100 m mitigation structure per 1 km length of infrastructure in critical tiger corridors (Wildlife institute of India 2016). Although we cannot test the effectiveness of different kinds of structures which can provide connectivity across roads in this simulation, we investigate the effect of having a gap in the barrier which can potentially maintain connectivity. |
| F8- Habitat restoration to establish corridors between all PAs | The corridors were designated based on the least cost paths (generated using the gdistance package(van Etten 2012) in R (Ihaka & Gentleman 2012)) between PAs and the proposed corridor between Kanha and Pench. | Restoration of habitat and establishing corridors between PAs has been recommended to maintain and even increase the connectivity in the landscape (Dinerstein *et al.* 2006; Dutta *et al.* 2015). We generated least cost paths between PAs and we assigned a value of 1 to all the pixels in these paths (equivalent to forest). All non-forest pixels within the least cost paths were converted to forest to represent restoration of habitat. These paths overlap with the corridors delineated in previous studies (Qureshi *et al.* 2014; Dutta *et al.* 2015). We test how beneficial establishing these corridors would be. |
| F9- Fenced PAs | All the protected areas have fence around them in the future preventing dispersal | Extreme scenario to investigate the effect of fencing on genetic variation and extinctions in the future. |

\* We carried out additional simulations for scenarios F1 and F4 where we added a 5 km buffer around the smaller PAs in the landscape (< 400 km<sup>2</sup>). We evaluated the effect of increased PA size on connectivity and extinction estimates.

The simulations included eleven additional unsampled PAs: PAN, S-U, Indravati Tiger Reserve (IND), Melghat Tiger Reserve (MEL), Ratapani Wildlife Sanctuary (RAT), Kawal Tiger Reserve (KAW), Tamorpingla Wildlife Sanctuary (TAM), Sanjay Tiger Reserve (SAN), Noradehi Wildlife Sanctuary (NOR), Barnawapara Wildlife Sanctuary (BAR)and Chaprala Wildlife Sanctuary (CHH). These areas were included in future scenario simulations as some have tigers, while others have high probability of tiger occupancy, or are protected reserves.

## Results

### Population Genetic Analysis

Out of 289 samples identified as tigers, data for at least 8 out of 12 microsatellite loci could be generated from 127 samples, 116 of which were identified as unique individuals. The P(ID) (the probability of two different individuals having the same genotype) was 1.48 x 10^−11^ and the more conservative measure Sib P(ID) (PID when all individuals in the population are assumed to be siblings) was 5.1 x 10^−5^, indicating that even related individuals would have a very low probability of having identical genotypes. The estimated allelic dropout across loci was 0.062 and the frequency of false alleles was 0.013 (comparable to other studies in the landscape; (Caragiulo *et al*. 2015)). Six out of the 12 loci showed significant deviation from Hardy-Weinberg equilibrium, suggesting the presence of genetic structure. The mean number of alleles per locus was 8.7 and expected heterozygosity was 0.723 (Supplementary Table S2). Global *F*_ST_ was estimated to be 0.169, with highest pairwise differentiation between BTR and BPR (Tables S2 and S3). Table 2 presents D_PS_, a measure of contemporary genetic differentiation.

**Table 2.** Genetic differentiation-DPS

|  | BTR<br>(n= 20) | BPR<br>(n= 18) | BOR<br>(n= 4) | KTR<br>(n= 22) | CHA<br>(n=8) | PTR<br>(n= 15) | STR<br>(n= 4) | TATR<br>(n= 26) | UK<br>(n= 5) |
| --- | --- | --- | --- | --- | --- | --- | --- | --- | --- |
| BTR | 0 |  |  |  |  |  |  |  |  |
| BPR | 0.629 | 0 |  |  |  |  |  |  |  |
| BOR | 0.669 | 0.456 | 0 |  |  |  |  |  |  |
| KTR | 0.548 | 0.597 | 0.641 | 0 |  |  |  |  |  |
| CHA | 0.621 | 0.286 | 0.359 | 0.592 | 0 |  |  |  |  |
| PTR | 0.501 | 0.627 | 0.612 | 0.351 | 0.637 | 0 |  |  |  |
| STR | 0.541 | 0.612 | 0.598 | 0.3541 | 0.592 | 0.416 | 0 |  |  |
| TATR | 0.639 | 0.213 | 0.413 | 0.571 | 0.267 | 0.612 | 0.591 | 0 |  |
| UK | 0.700 | 0.358 | 0.373 | 0.624 | 0.474 | 0.635 | 0.624 | 0.322 | 0 |

### Functional connectivity

Human settlements were the most important (highest magnitude of correlation) landscape variable explaining genetic distance between PAs. Land-use and traffic intensity on roads also explained significant variation, even after accounting for geographic distance (Table 3). These three variables (traffic intensity on roads, human settlements and land-use) were retained for multivariate optimization. The parameter estimates for each landscape variable are presented in Table 3.

**Table 3.**
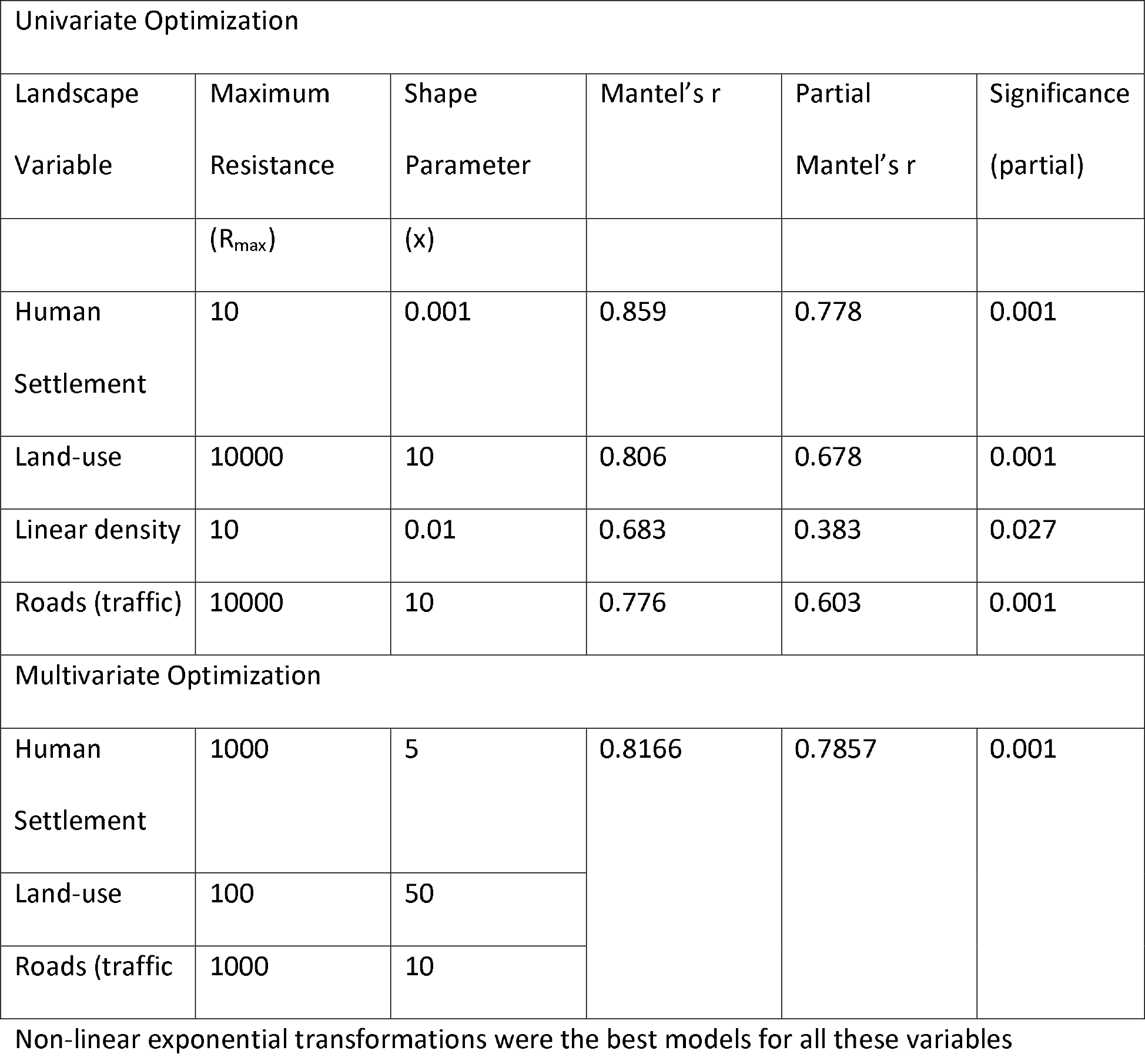
Univariate optimization results

| Univariate Optimization |  |  |  |  |  |
| --- | --- | --- | --- | --- | --- |
| Landscape<br>Variable | Maximum<br>Resistance | Shape<br>Parameter | Mantel's r | Partial<br>Mantel's r | Significance<br>(partial) |

| | ($R_{\max}$) | (x) | | | |
| --- | --- | --- | --- | --- | --- |
| Human Settlement | 10 | 0.001 | 0.859 | 0.778 | 0.001 |
| Land-use | 10000 | 10 | 0.806 | 0.678 | 0.001 |
| Linear density | 10 | 0.01 | 0.683 | 0.383 | 0.027 |
| Roads (traffic) | 10000 | 10 | 0.776 | 0.603 | 0.001 |
| Multivariate Optimization |  |  |  |  |  |
| Human Settlement | 1000 | 5 | 0.8166 | 0.7857 | 0.001 |
| Land-use | 100 | 50 |  |  |  |
| Roads (traffic) | 1000 | 10 |  |  |  |

Non-linear exponential transformations were the best models for all these variables

Shape parameter (the parameter that determines the shape of the relationship between the landscape variable and resistance) and maximum resistance of the optimum models of all the three landscape variables changed on combining, suggesting interaction between these variables. The non-linear transformations (x >1) indicate that low resistance is offered by smaller and middle values assigned to the variable and the resistance increases steeply with very high values. For example, roads with low and moderate traffic offer negligible resistance to movement, however, the resistance increases steeply with very high traffic. Correlation between the pairwise cost distance among populations (estimated from the combined resistance surface) and genetic distance was high (0.7857 after controlling for the geographic distance, 0.8166 without controlling for geographic distance). The correlation value for isolation by distance model (geographical distance alone) was 0.624

### Future connectivity

Overall, genetic diversity reduced over time in all the simulation scenarios (Figure S6). However, restoring and protecting corridors between PAs lead to the least decline in genetic variation (~20%). Heterozygosity decreased faster and reduced to a lower final value at the end of 100 years in the scenarios with lower dispersal threshold (300km). Within both the dispersal categories, the loss of genetic diversity was greater in the scenarios where the forest cover loss was higher (Figure S6). Our results suggest that inbreeding did not increase appreciably over 100 years (F<0.25 in all scenarios). Figure 2 summarizes the implications of three different management decisions on structural connectivity, allelic richness, inbreeding and extinction in a subset of the simulated scenarios.

**Figure 2.**
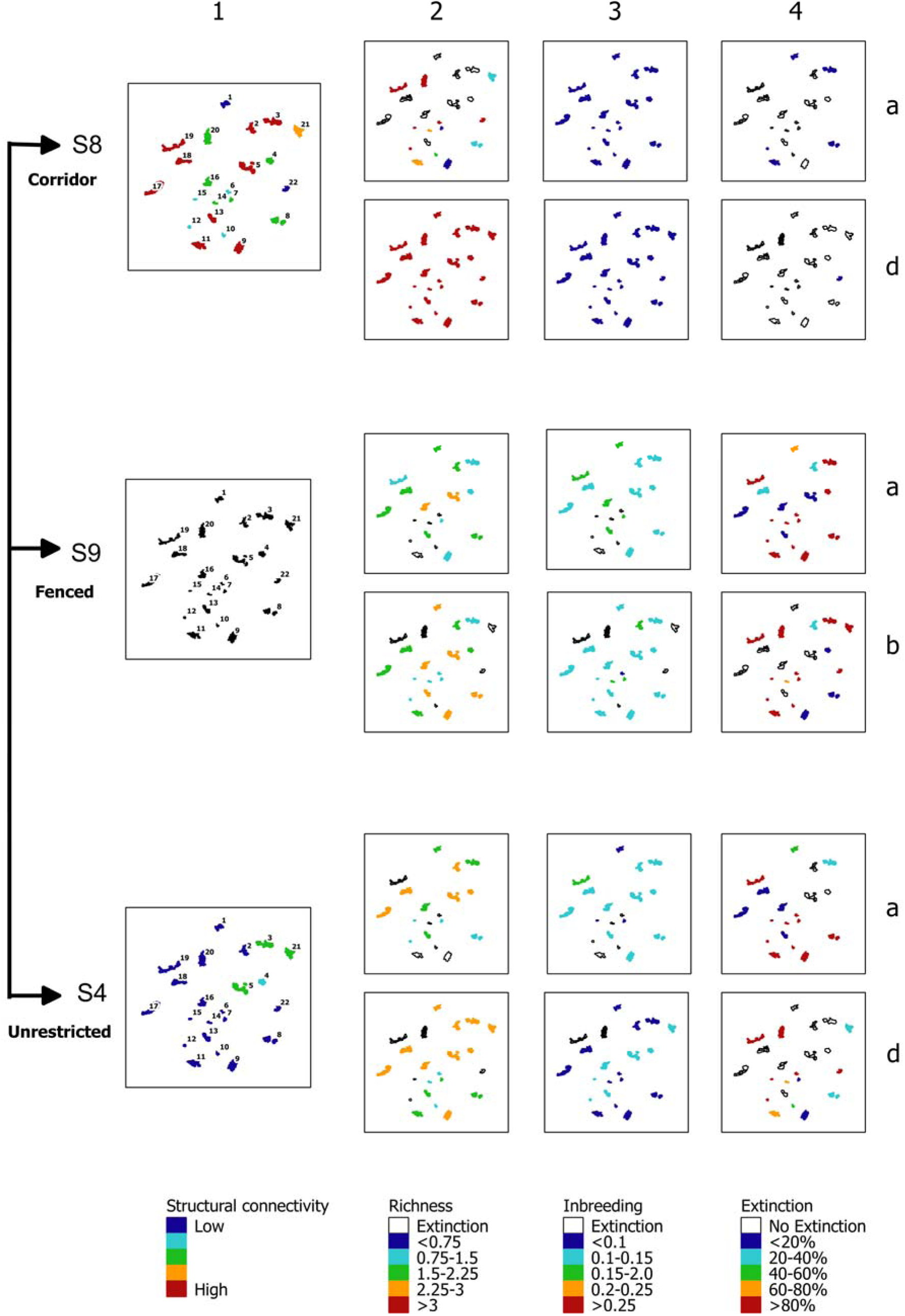
Structural connectivity, allelic richness, inbreeding and extinction after 100 years under selected management scenarios. The 3 management scenarios in this figure have corresponding panels with two sub-scenarios (a- tiger number does not increase and d- increase in tiger number) and 4 plots each representing management outcomes for (L to R): 1-structural connectivity, 2-allelic richness, 3-inbreeding estimate, and 4-extinction probability. The structural connectivity is the same for sub-scenarios a and d. Hence there is a single structural connectivity plot for each of the management outcomes. Completely isolated PAs (scenario F9) are represented in black. The outcomes for different scenarios can be compared within a column. The coloured areas in each plot are the protected areas and their colour represents the scenario-specific outcome. The gradient from blue to red represents low to high (structural connectivity, allelic richness, inbreeding coefficient and extinction probability). Blank PAs in the plots within columns 2 and 3 are the PAs which go extinct by the year 2100. Blank PAs in column 4 (extinction probability plots) are the ones which do not show extinction in any of the 100 replicate simulation runs.

Irrespective of land-use change scenario, dispersal threshold and tiger demographic trajectory, small isolated PAs (TIP and BOR) had the highest risk of extinction. Reducing the dispersal threshold from 500km to 300km doubled the average extinction probability of some populations (BTR, PTR, MEL and STR, see Figure 3a and 3b). Currently well connected, but small PAs (UK, CHH, NGZ and NAW) had high extinction probability only in the scenarios where forest cover around them was lost. Some large PAs that currently have a very low number of tigers (< 10 tigers, KAW, IND, RAT and S-U) also had high extinction probability except in the sub-scenarios where tiger numbers increased. Small population size was associated with high variance and highly skewed sex ratios (Figures S7 and S8). The extinction probability of small PAs was also governed by their isolation and associated re-colonization probability (Figure 3). Adding a buffer around the small PAs reduced their overall extinction probability. Among the isolated small PAs (TIP and BOR), addition of buffer reduced the extinction probability by ~23%. Adding a buffer zone around the small, currently connected populations reduced the extinction probability to a large extent (~70%), but only in the simulations where the landscape around the buffers changed (F4). The benefit of having a buffer around the small PAs was the highest in the sub-scenario with stepping stone populations (sub-scenario c) between PAs.

**Figure 3.**
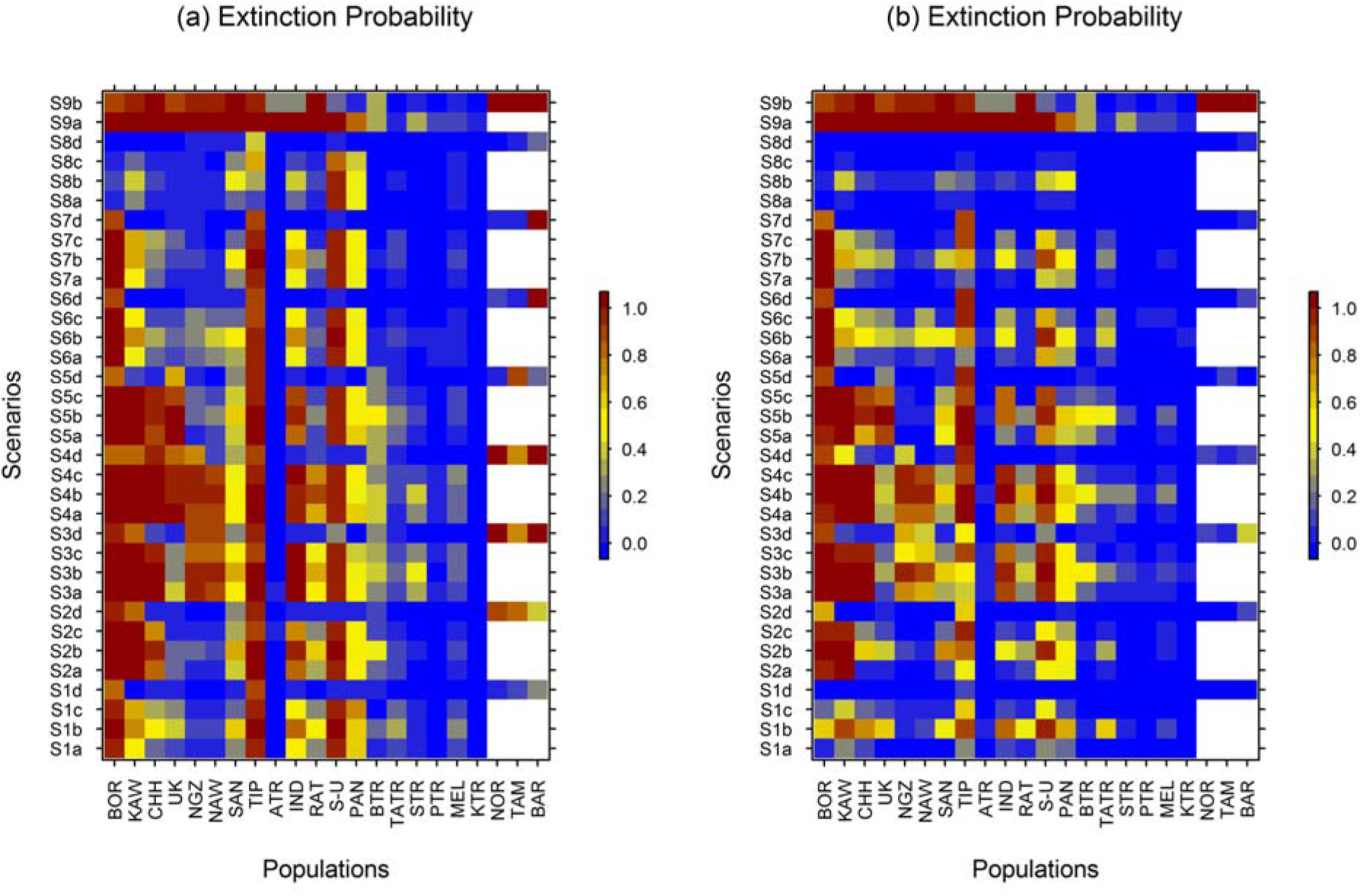
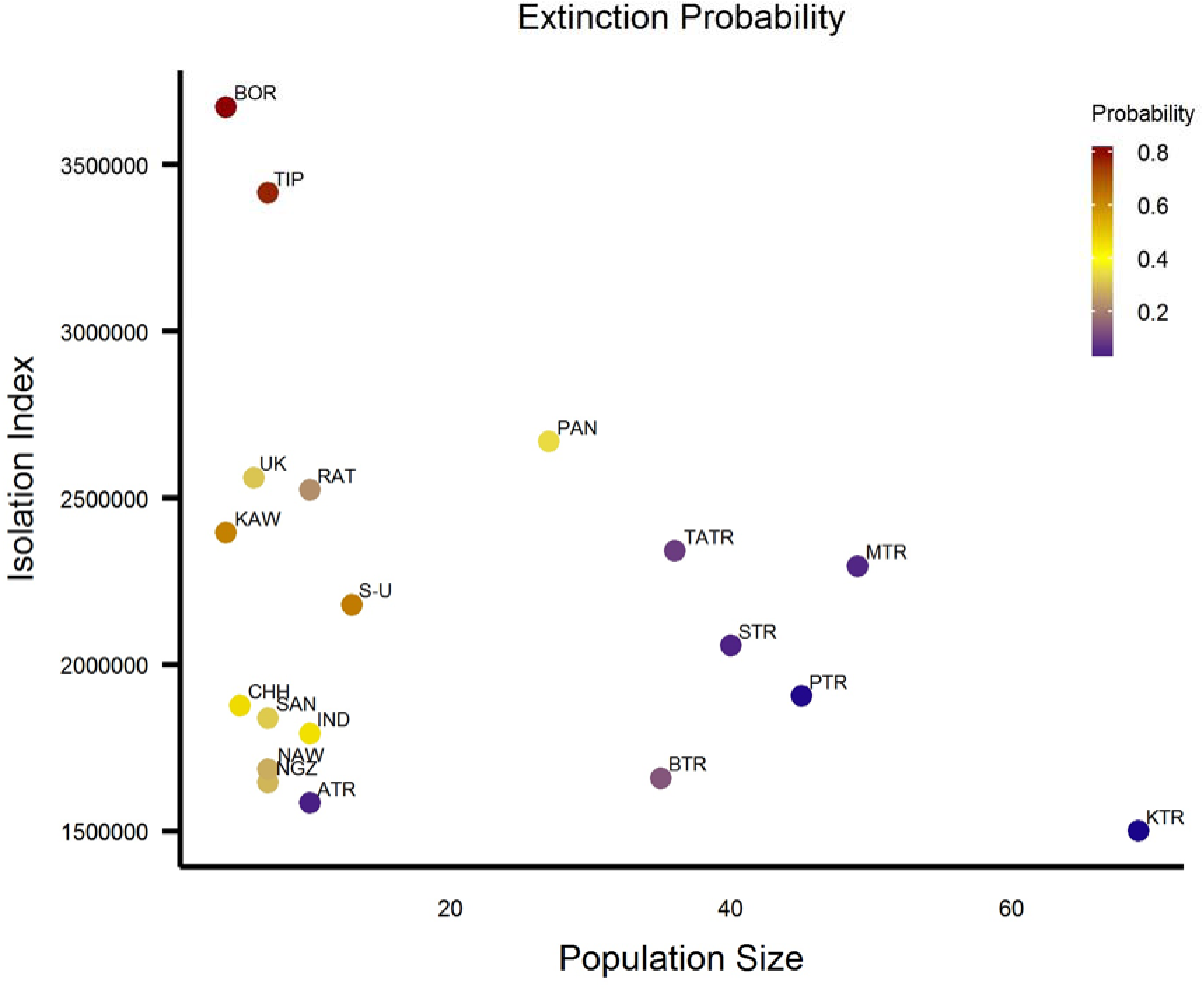
Extinction Probability. (a) Matrix representing extinction probability for each population for each of the scenarios after 100 yearsdispersal threshold 300km (b) Matrix representing extinction probability for each population for each of the scenarios after 100 years- dispersal threshold 500km. x-axis represents the PAs and y axis represents the scenarios and sub-scenarios (c) Scatterplot of population size (average of current size and size in sub-scenario d) vs. isolation index calculated as the average cost distance between populations. Each point represents a population and the colour represents it’s average extinction probabilityacross scenarios. NOR, TAM, BAR do not have any tigers currently and were only included in the simulation sub-scenario d where tiger numbers increase. Hence they do not have colour in panels (a) and (b) for scenarios a, b and c. Scenarios: F1-No change in landscape, F2-Forest cover constant, F3-Area under agriculture constant, F4-Unconstrained landscape change, F5-Mines, F6-NH6 and NH7 as barriers, F7-NH6 and NH7 as barriers with gaps, F8-Corridors, F9-PAs fenced. Sub-scenarios: a- Tigers only inside protected areas, b- Tigers inside and outside (random), c- Tigers inside and outside (clustered), d- Tiger numbers increase.

### Change in connectivity: specific infrastructure projects

Increase in mined area and associated increase in built-up areas lead to ~15 times higher extinction probability in small and medium sized PAs of BTR, SAN, UK and KAW due to their proximity to coal fields. Presence of NH7 as a barrier without any mitigation structures (scenario F6) increased the *F*_ST_ between KTR and PTR ~4 times compared to scenario F1 and scenario F7 (Figure S9). NH7 bisects the corridor between these two PAs. NH6 bisects the corridor between NGZ and NAW. The scenario with NH6 as a barrier (scenario F6) leads to ~19 times and ~65 times higher probability of extinction for dispersal threshold of 500 km and 300 km respectively. Higher increase in the genetic differentiation in the latter case (between NGZ-NAW, as compared to KTR-PTR) could be due to the smaller current population sizes of NGZ and NAW. Across land-use change scenarios, increasing tiger number (sub-scenario d) within and outside PAs (sub-scenarios c) lead to ~37% lower extinction probabilities overall and ~11% lower reduction in heterozygosity.

## Discussion

The St. Petersburg declaration of 2010 envisaged doubling tiger numbers across all tiger nations by 2022 (Global Tiger Recovery Program 2011). Our results from CIL demonstrate that maintaining or establishing connectivity and ensuring protection will be critical to meet such tiger conservation targets. Along with corridor conservation, designing, notifying and maintaining stepping-stone populations between PAs will be beneficial for maintaining future connectivity. To facilitate regional planning, spatially explicit analysis could help identify locations where infrastructure could be developed without affecting tiger connectivity. The current results are general prescriptions, but findings from this study provide evidence that can be translated into spatially explicit conservation planning exercises. Additionally, our framework can be applied globally to other landscapes and species (where genetic data are available) to improve connectivity in the context of development.

### What impacts current tiger connectivity?

Human footprint on the landscape had the strongest impact on connectivity. Dense human settlements and roads with high traffic offered highest resistance to movement. Degraded forests and plantations offered negligible resistance and agriculture-village matrix offered low resistance to tiger movement. However, long contiguous stretches of agriculture- village matrix could lead to accumulation of cost over space and impede movement. Our results are supported by empirical data on tiger movement. Recent data from radio-collared tigers reveals that long distance dispersing tigers do not avoid agriculture-village matrix and cross low traffic roads (Athreya *et al*. 2014; Krishnamurthy *et al*. 2016). Both, traffic intensity and dense human settlements have a non-linear relationship with resistance, suggesting that only very high intensity traffic and high-density human settlements offered high resistance to movement (Table 4).

Roads are known to negatively impact genetic diversity and differentiation in animal species, especially for mammals and amphibians (Shirk *et al*. 2010; Holderegger & Di Giulio 2010). However, most of these studies have been carried out in developed/industrialized countries where road density is high and many of the roads are fenced (Holderegger & Di Giulio 2010). Although India has the second largest road network in the world, 46% of the roads are currently not surfaced (National Transport Policy Development Committee 2013)and very few segments are fenced. As a result, even the existing national highways connecting major centers may be permeable to tiger movement on stretches with low traffic volumes and when other landscape features promote dispersal. However, this is bound to change in the future as roads will be widened to accommodate increasing traffic volumes.

### Future change in diversity

Genetic diversity reduced over time in all simulated scenarios. Results suggest that even establishing corridors would not suffice to maintain current level of heterozygosity. Our results are supported by Bay *et al*. (2013), where simulations of mitochondrial diversity revealed that even with geneflow, the number of tigers essential to maintain current heterozygosity would exceed the carrying capacity of PAs in peninsular India (Bay *et al*. 2013). Our simulations revealed that both increasing tiger numbers and maintaining connectivity were essential for preventing drastic reduction of heterozygosity.

### Stepping-stone corridors preserve connectivity

Our simulations revealed that loss of forest cover due to diversion of land for agriculture or infrastructure lead to high genetic differentiation. However, increasing number of tigers within existing PAs and allowing clusters of individuals to survive outside PAs decreased the observed genetic differentiation and inbreeding. Presence of breeding clusters of tigers outside PAs also reduced the probability of extinction. These intervening clusters aid in dispersal between the larger, more robust populations, thus forming ‘stepping-stone corridors’. Scenarios that incorporated stepping-stone corridors resulted in 10% higher genetic variation and 6-86% lower extinction probability compared to those without.

### Habitat restoration and protection is critical

Dinerstein et al. (2006) recommended restoring habitat to increase population connectivity between tiger conservation landscapes. Our results demonstrated that such habitat restoration to establish corridors reduced extinction probability by 68% (Scenario F8, Figure 3a and 3b) and was critical for population persistence in the future. Such landscape restoration may require careful selection of areas so as to benefit both people and wildlife (Defries *et al*. 2007). This may be difficult to achieve between all pairs of PAs, especially those that have negligible structural connectivity between them. Our results revealed that increasing tiger numbers allowed large but currently degraded PAs (with few tigers) to achieve low extinction probability, underscoring the importance of better PA management and protection. Increased tiger numbers buffered against demographic stochasticity in small PAs and decreased the overall extinction probability. This was best demonstrated in the scenario where we added a buffer around the small PAs and extinction probability dropped by 23-70%. The government of India is taking steps to notify ‘Eco-sensitive Zones’ around PAs to regulate developmental activities (Mathur 2012). However, our results suggest that unless connectivity is restored and stepping-stone populations protected (also suggested by (Chundawat *et al*. 2016)), even such rescue effects would fail for small PAs when the landscape around them became unsuitable. Such extinction debt poses a significant challenge for conservation while these tiger populations still persist.

### Low levels of inbreeding in Central Indian tigers of the future

Kenney *et al*. (2014) investigated the effect of inbreeding depression on population viability in tigers and found that even populations as large as (with 63-80 > 1 year old tigers) the big populations in our study have a high future risk of extinction due to inbreeding depression if connectivity was not maintained. Our results suggest that inbreeding did not increase appreciably over the next 100 years (F<0.25 in all scenarios). Levels of inbreeding were lower than those known to impact fitness in mammals based on studies in the wild and captivity (Ralls & Ballou 1982; Keller 2002). Hence, we did not simulate the effect of inbreeding depression on survival and extinction. However, further increase in the inbreeding co-efficient over time may lead to inbreeding depression and increase extinction risk of even large populations. Our simulations suggest that inbreeding coefficient is likely to increase faster if tiger numbers do not increase in the future (Figure S12).

### Model assumptions and caveats

Overall, our framework provides the first snapshot of the potential to combine landscape genetics with spatially explicit simulations for tiger conservation. Future studies should investigate how different land-use change data and dispersal strategies would affect tiger connectivity in CIL. For example, future landscape change projection in our simulations was based on landscape change over one decade (2001-2012). Would longer decadal datasets help us better understand LULC change in the CIL? Additionally, our model does not allow for future land-use policy changes. For example, accelerated development could result in loss of connectivity in less than 100 years. However, we do incorporate scenarios which demonstrate the impacts of specific infrastructure projects currently being undertaken or envisaged in the future (Scenarios F5-F7).

While modeling tiger dispersal, we did not consider the variation in behavioral response of individuals to land use change and the probability of locating a corridor. Modeling how individuals incorporate information about surrounding habitat while dispersing would improve the simulation results (Colbert *et al*. 2009). However, such data is currently not available. Future research should evaluate the influence of different dispersal strategies on tiger connectivity. Finally, our simulations investigating tiger connectivity assume a maximum dispersal distance, which is difficult to estimate in the wild. The two maximum dispersal distances of 300 and 500km that we used were based on available data (Bowman *et al*. 2002; Patil *et al*. 2011; Natesh *et al*. 2017). A possibility of longer than 500 km dispersal cannot be ruled out, and hence we carried out simulations with a longer maximum dispersal distance (as suggested in (Joshi *et al*. 2013), results in supplementary material). This longer threshold led to lower genetic differentiation (in scenario F1) than currently observed, suggesting that this particular dispersal scenario may be overestimating the probability of long distance dispersals. However, even with such enhanced dispersal distances, the key populations that needing management attention, the impact of infrastructure projects, and scenarios that lead to better outcomes did not change (Supplementary Figures S6, S9-S12).

### Implications for Conservation Planning

Our results reveal that multiple actions like delineating and maintaining corridors, with stepping-stone populations between PAs, along with increasing tiger numbers are necessary to maintain long-term viable tiger populations at a landscape scale. Our results highlight the immediate need for regional land-use management and planning exercises aimed at managing tiger populations as a network of PAs connected with corridors.

### Land-use change

Currently, infrastructure development does not incorporate conservation goals while developing project plans, which could seriously undermine ongoing tiger conservation efforts within the landscape. Nearly 50% of India’s population is projected to live in cities by 2030 (World Bank Group 2015), and research has shown that built- up area increases faster than population increase in urban areas (Sudhira *et al*. 2004). Along with urbanization, coal requirement for electricity generation is projected to increase ~2.5 times by 2031-32 (Fernandes 2012). To provide better connectivity between cities and to accommodate increasing road traffic (estimated to grow at about 13% per year over the next 20 years), massive infrastructure development projects are being undertaken (National Transport Policy Development Committee 2013). Our simulations reveal the impacts of upcoming development on tiger connectivity. Ideally, expansion of the current road network should include realignment of new roads to avoid critical tiger habitat (see (Raman 2011)). We show that widening national highways without installing crossing structures (NH6 and NH7, Scenario F6) in an area critical to tiger connectivity will increase genetic differentiation between populations on either side 19 to 65 fold. When alternative routes are not available, we strongly recommend planning and installing mitigation structures (under and over-passes) for wildlife passage before roads are built or widened. This is more economical than retrofitting existing roads and should be considered during the environment impact assessment of new infrastructure projects (Glista *et al*. 2009). Our results (Figure S9) highlight the importance of installing crossing structures for maintaining connectivity (F_ST_ increases by 80% from an average 0.025 to 0.13 in the absence of such structures). Currently, such regional level planning which aligns conservation goals with developmental plans is in its infancy in India. Our results provide impetus to such efforts by highlighting landscape variables that need to be considered while developing infrastructure plans.

### Forest diversion and mining

Diversion of forest for mining is one of the major causes for loss of forest cover and structural connectivity within the CIL. Over the last 3 decades, 40% of the total forest land diverted has been for mining (Centre for Science and Environment 2012). Coal mining alone accounted for 65% of the total land diverted for mining between 2007 and 2011 (Centre for Science and Environment 2012). If, along with other minerals being mined, all the coal blocks in the study landscape were opened for mining, the mined area and associated increase in built- up area would lead to ~22 % higher extinction probability even for large PAs in proximity to these blocks. We emphasize the urgent need to protect the delineated corridors (as in Qureshi *et al*. 2014) for preserving connectivity to prevent extinctions.

## Conclusion

Several species globally face threats due to anthropogenic impacts, similar to the tigers in our study area. Landscapes with high conservation value, especially in the tropics, have been changing rapidly due to increasing human population and exploitation of natural resource leading to conversion and degradation of habitat (Crooks *et al*. 2011; Elmhagen *et al*. 2015; Newbold *et al*. 2015). Our results highlight the urgent need for informed development plans that consider biodiversity and connected wildlife populations in addition to human development goals. With habitats of most large mammals getting increasingly fragmented, our approach of combining landscape genetics with forward-time simulations to estimate extinction probability and loss of connectivity provides a valuable tool for conservation management. Such an approach can help identify populations vulnerable to landscape change, improving conservation and management of populations, species and landscapes to ensure long-term persistence.

## Acknowledgements

We are thankful to the Maharashtra, Madhya Pradesh and Chhattisgarh Forest Departments for providing the necessary permissions and logistical support for this research. We sincerely thank the forest department staff, field assistants and volunteers for their help and support during the fieldwork. We thank the Indian Resource Center for Clark Labs at Ashoka Trust for Research in Ecology and the Environment for letting us use their facilities. We thank V. V. Robin, Meghana Natesh, Y. V. Jhala and an anonymous reviewer for helpful comments on the manuscript. We are grateful to the National Tiger Conservation Authority (NTCA) of India for partially funding this study. We thank the National Centre for Biological Sciences (NCBS) for institutional support. PT was supported by Council for Industrial and Scientific Research fellowship and NCBS. UR was supported by the DAE outstanding Scientist Award.

## References

Adamack AT, Gruber B (2014) PopGenReport: simplifying basic population genetic analyses in R (S Dray, Ed,). Methods in Ecology and Evolution, 5, 384–387.

Areendran G, Rao P, Raj K, Mazumdar S, Puri K (2013) Land use / land cover change dynamics analysis in mining areas of Singrauli district in Madhya Pradesh, India. Tropical Ecology, 54, 239–250.

Athreya V, Navya R, Punjabi G a et al. (2014) Movement and activity pattern of a collared tigress in a human-dominated landscape in central India. Tropical Conservation Science, 7, 75–86.

Balloux F (2001) EASYPOP (Version 1.7): A Computer Program for Population Genetics Simulations. Journal of Heredity, 92, 301–302.

Barnosky AD, Matzke N, Tomiya S et al. (2011) Has the Earth’s sixth mass extinction already arrived? Nature, 471, 51–7.

Bay RA, Ramakrishnan U, Hadly EA (2013) A call for tiger management using “reserves” of genetic diversity. The Journal of heredity, 105, 295–302.

Benson JF, Mahoney PJ, Sikich JA et al. (2016) Interactions between demography, genetics, and landscape connectivity increase extinction probability for a small population of large carnivores in a major metropolitan area. Proceedings of the Royal Society of London B: Biological Sciences, 283.

Boersen MR, Clark JD, King TL (2003) Estimating black bear population density and genetic diversity at Tensas River, Louisiana using markers microsatellite. Wildlife Society Bulletin, 31, 197–207.

Bowcock AM, Ruiz-Linares A, Tomfohrde J et al. (1994) High resolution of human evolutionary trees with polymorphic microsatellites. Nature, 368, 455–7.

Bowman J, Jaeger J a. G, Fahrig L (2002) Dispersal Distance of Mammals Is Proportional To Home Range Size. Ecology, 83, 2049–2055.

Brown JL, Weber JJ, Alvarado-Serrano DF et al. (2016) Predicting the genetic consequences of future climate change: The power of coupling spatial demography, the coalescent, and historical landscape changes. American journal of botany, 103, 153–63.

Caragiulo A, Pickles RSA, Smith JA et al. (2015) Tiger (Panthera tigris) scent DNA: a valuable conservation tool for individual identification and population monitoring. Conservation Genetics Resources, 7, 681–683.

Centre for Science and Environment (2012) Forest and Environment Clearances. Centre for Science and Environment, New Delhi, India.

Chundawat RS, Sharma K, Gogate N, Malik PK, Tamim A (2016) Size matters: Scale mismatch between space use patterns of tigers and protected area size in a Tropical Dry Forest. Biological Conservation, 197, 146–153.

Colbert J, Le Galliard JF, Cote J, Meylan S, Massot M (2009) Informed dispersal, heterogeneity in animal dispersal syndromes and the dynamics of spatially structured populations. Ecology Letters, 12, 197–209.

Crooks KR, Burdett CL, Theobald DM, Rondinini C, Boitani L (2011) Global patterns of fragmentation and connectivity of mammalian carnivore habitat. Philosophical transactions of the Royal Society of London. Series B, Biological sciences, 366, 2642–51.

Defries RS, Hansen A, Turner BL, Reid R, Liu J (2007) Land use change around protected areas: management to balance human needs and ecological function. Ecological Applications, 17, 1031–1038.

Dieringer D, Schlotterer C (2003) microsatellite analyser (MSA): a platform independent analysis tool for large microsatellite data sets. Molecular Ecology Notes, 3, 167–169.

Dinerstein E, Loucks C, Heydlauff A et al. (2006) Setting Priorities for the Conservation and Recovery of Wild Tigers: 2005–2015. A User’s Guide. WWF, WCS, Smithsonian, and NFWF-STF, Washington, D.C. – New York.

Dirzo R, Young HS, Galetti M et al. (2014) Defaunation in the Anthropocene. Science, 345, 401–406.

Dooley MP, Folkerts-landau D, Garber PM (2014) The revived Bretton Woods system’s first decade. National Bureau of Economic Research, Cambridge.

Dutta T, Sharma S, Mcrae BH, Sarathi P, Defries R (2015) Connecting the dots⍰: mapping habitat connectivity for tigers in central India. Regional Environmental Change.

Elmhagen B, Eriksson O, Lindborg R (2015) Implications of climate and land-use change for landscape processes, biodiversity, ecosystem services, and governance. Ambio, 44, S1–S5.

van Etten J (2012) R package gdistance: distances and routes on geographical grids (version 1.1-1).

Fernandes A (2012) How coal mining is thrashing tigerland. Greenpeace, India.

Forest Survey of India (2015) India State of Forest Report 2015. Forest Survey of India, Dehradun, India.

Frankham R (1996) Relationship of Genetic Variation to Population Size in Wildlife. Conservation Biology, 10, 1500–1508.

Glista DJ, DeVault TL, DeWoody JA (2009) A review of mitigation measures for reducing wildlife mortality on roadways. Landscape and Urban Planning, 91, 1–7.

Global Tiger Recovery Program (2011) Global Tiger Recovery Program 2010–2022. The World Bank, Washington, DC.

Goudet J (2005) hierfstat, a package for r to compute and test hierarchical F-statistics. Molecular Ecology Notes, 5, 184–186.

Hagen SB, Kopatz A, Aspi J, Kojola I, Eiken HG (2015) Evidence of rapid change in genetic structure and diversity during range expansion in a recovering large terrestrial carnivore. Proceedings. Biological sciences / The Royal Society, 282, 20150092.

Holderegger R, Di Giulio M (2010) The genetic effects of roads: A review of empirical evidence. Basic and Applied Ecology, 11, 522–531.

Ihaka R, Gentleman R (2012) R: A Language for Data Analysis and Graphics. Journal of Computational and Graphical Statistics.

Jackson CR, Marnewick K, Lindsey PA, Røskaft E, Robertson MP (2016) Evaluating habitat connectivity methodologies: a case study with endangered African wild dogs in South Africa. Landscape Ecology, 31, 1433–1447.

Jhala, Y. V.; Qureshi, Q.; Gopal R. (2015) Status of tigers in India, 2014. National Tiger Conservation Authority, New Delhi & The Wildlife Institute of India, Dehradun.

Jombart T (2008) adegenet: a R package for the multivariate analysis of genetic markers. Bioinformatics (Oxford, England), 24, 1403–5.

Joshi A, Vaidyanathan S, Mondol S, Edgaonkar A, Ramakrishnan U (2013) Connectivity of Tiger (Panthera tigris) Populations in the Human-Influenced Forest Mosaic of Central India. PLoS ONE, 8, e77980.

Keller L (2002) Inbreeding effects in wild populations. Trends in Ecology & Evolution, 17, 230–241.

Krishnamurthy R, Cushman SA, Sarkar MS et al. (2016) Multi-scale prediction of landscape resistance for tiger dispersal in central India. Landscape Ecology, 31, 1355–1368.

Landguth EL, Muhlfeld CC, Waples RS et al. (2014) Combining demographic and genetic factors to assess population vulnerability in stream species. Ecological Applications, 24, 1505–1524.

Laurance WF, Balmford a. (2013) A global map for road building. Nature, 495, 308–309.

Lele S, Rosencranz A (2008) Supreme Court and India’s Forests. Economic and Political Weekly, 11–14.

Mathur VB (2012) India’s action plan for implementing the convention on biological diversity’s programme of work on protected areas. Wildlife Institute of India, Dehradun; Ministry of Environment and Forests, New Delhi, India.

Ministry of Environment and Forests (2008) National Action Plan on Climate Change. Ministry of Enivronment and Forests, New Delhi, India.

Ministry of Mines (2011) Unlocking the Potential of the Indian Minerals Sector. Strategy Paper for Ministry of Mines, New Delhi, India.

Mondol S, Karanth KU, Ramakrishnan U (2009a) Why the Indian subcontinent holds the key to global tiger recovery. PLoS genetics, 5, e1000585.

Mondol S, Ullas Karanth K, Samba Kumar N et al. (2009b) Evaluation of non-invasive genetic sampling methods for estimating tiger population size. Biological Conservation, 142, 2350–2360.

Morrison JC, Sechrest W, Dinerstein E, Wilcove DS, Lamoreux JF (2007) Persistence of Large Mammal Faunas as Indicators of Global Human Impacts. Journal of Mammalogy, 88, 1363–1380.

Mukherjee N, Mondol S, Andheria A, Ramakrishnan U (2007) Rapid multiplex PCR based species identification of wild tigers using non-invasive samples. Conservation Genetics, 8, 1465–1470.

Natesh M, Alta G, Nigam P et al. (2017) Conservation priorities for endangered Indian tigers through a genomic lens. Scientific Reports, 7.

National Transport Policy Development Committee (2013) Trends in Growth and development of transport. Planning Commission, India.

Newbold T, Hudson LN, Hill SLL et al. (2015) Global effects of land use on local terrestrial biodiversity. Nature, 520, 45–50.

Patil N, Samba Kumar N, Gopalaswamy AM, Ullas Karanth K (2011) Dispersing tiger makes a point. Oryx, 45, 472–475.

Prakash A, Gupta RP (2016) Land-use mapping and change detection in a coal mining area - a case study in the Jharia coalfield, India. International Journal of Remote Sensing, 19, 391–410.

Qureshi Q, Saini S, Basu P et al. (2014) Connecting tiger populations for long-term conservation. National Consevation Authority of India & Wildlife Institute of India, Dehradun. TR2014-02.

Ralls K, Ballou J (1982) Effect og inbreeding on juvenile mortality in some small mammal species. Laboratory Animals, 16, 159–166.

Raman TRS (2011) Framing ecologically sound policy on linear intrusions affecting wildlife habitats: Background paper for the national board of wildlife. Nature Conservation Foundation, Mysore.

Reddy PA, Cushman SA, Srivastava A, Sarkar MS, Shivaji S (2017) Tiger abundance and gene flow in Central India are driven by disparate combinations of topography and land cover. Diversity and Distributions, 1–12.

Ripple WJ, Beschta RL (2012) Trophic cascades in Yellowstone: The first 15years after wolf reintroduction. Biological Conservation, 145, 205–213.

Ripple WJ, Estes JA, Beschta RL et al. (2014) Status and ecological effects of the world’s largest carnivores. Science, 343, 1241484.

Saccheri I, Kuussaari M, Kankare M, Vikman P, Hanski I (1998) Inbreeding and extinction in a butterfly metapopulation. Nature, 392, 491–494.

Sharma S, Dutta T, Maldonado JE et al. (2013a) Selection of microsatellite loci for genetic monitoring of sloth bears. Ursus, 24, 164–169.

Sharma S, Dutta T, Maldonado JE et al. (2013b) Forest corridors maintain historical gene flow ina tiger metapopulation in the highlands of central India. Proceedings of the Royal Society B: Biological Sciences, 280, 20131506.

Shirk AJ, Wallin DO, Cushman SA, Rice CG, Warheit KI (2010) Inferring landscape effects on gene flow: a new model selection framework. Molecular Ecology, 19, 3603–3619.

Smouse PE, Long JC, Sokal RR (1986) Multiple Regression and Correlation Extensions of the Mantel Test of Matrix Correspondence. Systematic Zoology, 35, 627.

Spielman D, Brook BW, Briscoe DA, Frankham R (2004) Does Inbreeding and Loss of Genetic Diversity Decrease Disease Resistance? Conservation Genetics, 5, 439–448.

Srivastava R, Tyagi R (2016) Wildlife corridors in india. Environmental Law Review, 18, 205–223.

Sudhira HS (2011) Urban landcover and landcover change dataset of Indian cities. Indian Institute of Human Settlements Working Paper, Mimeo.

Sudhira HSS, Ramachandra TV V., Jagadish KSS (2004) Urban sprawl: metrics, dynamics and modelling using GIS. International Journal of Applied Earth Observation and Geoinformation, 5, 29–39.

Sutherland GD, Harestad AS, Price K, Lertzman KP (2000) Scaling of Natal Dispersal Distances in Terrestrial Birds and Mammals. Conservation Ecology, 4, 16.

Tian Y, Wu J, Smith AT et al. (2011) Population viability of the Siberian Tiger in a changing landscape: Going, going and gone? Ecological Modelling, 222, 3166–3180.

Tian Y, Wu J, Wang T, Ge J (2014) Climate change and landscape fragmentation jeopardize the population viability of the Siberian tiger (Panthera tigris altaica). Landscape Ecology, 29, 621–637.

Trinkel M, Cooper D, Packer C, Slotow R (2011) Inbreeding depression increases susceptibility to bovine tuberculosis in lions: an experimental test using an inbred-outbred contrast through translocation. Journal of wildlife diseases, 47, 494–500.

United Nations, Department of Economic and Social Affairs, Population Division (2014) World Urbanization Prospects: The 2014 Revision, Highlights (ST/ESA/SER.A/352).

Wikramanayake E, Dinerstein E, Loucks C et al. (2010) Roads to Recovery or Catastrophic Loss⍰: How Will the Next Decade End for Wild Tigers? (R Tilson, P Nyhus, Eds,). Elsevier Inc.

Wildlife institute of India (2016) Eco-friendly measures to mitigate impacts of linear infrastructure on wildlife. Wildlife Insitute of India, Dehradun, India.

Winter DJ (2012) MMOD: an R library for the calculation of population differentiation statistics. Molecular ecology resources, 12, 1158–60.

World Bank Group (2015) Population Estimates and Projections. The World Bank, Washington, DC.

Yadav PK, Kapoor M, Sarma K (2012) Land Use Land Cover Mapping, Change Detection and Conflict Analysis of Nagzira-Navegaon Corridor, Central India Using Geospatial Technology. International Journal of Remote Sensing and GIS, 1, 90–98.

Yumnam B, Jhala Y V, Qureshi Q et al. (2014) Prioritizing tiger conservation through landscape genetics and habitat linkages. PloS one, 9, e111207.

